# Leveraging a new data resource to define the response of *C. neoformans* to environmental signals

**DOI:** 10.1101/2023.04.19.537239

**Authors:** Yu Sung Kang, Jeffery Jung, Holly Brown, Chase Mateusiak, Tamara L. Doering, Michael R. Brent

**Author notes:** These authors contributed equally (Yu Sung Kang, Jeffery Jung). Corresponding author Michael Brent, http://mblab.wustl.edu, Campus Box 8510, Washington University, Saint Louis, MO 63130.

## Abstract

*Cryptococcus neoformans* is an opportunistic fungal pathogen with a polysaccharide capsule that becomes greatly enlarged in the mammalian host and during *in vitro* growth under host-like conditions. To understand how individual environmental signals affect capsule size and gene expression, we grew cells in all combinations of five signals implicated in capsule size and systematically measured cell and capsule sizes. We also sampled these cultures over time and performed RNA-Seq in quadruplicate, yielding 881 RNA-Seq samples. Analysis of the resulting data sets showed that capsule induction in tissue culture medium, typically used to represent host-like conditions, requires the presence of either CO_2_ or exogenous cyclic AMP (cAMP). Surprisingly, adding either of these pushes overall gene expression in the opposite direction from tissue culture media alone, even though both are required for capsule development. Another unexpected finding was that rich medium blocks capsule growth completely. Statistical analysis further revealed many genes whose expression is associated with capsule thickness; deletion of one of these significantly reduced capsule size. Beyond illuminating capsule induction, our massive, uniformly collected dataset will be a significant resource for the research community.

**IMPORTANCE:** *Cryptococcus neoformans* is an opportunistic yeast that kills ∼150,000 people each year. This major impact on human health makes it imperative to understand the basic biology of *C. neoformans* and the factors that mediate its virulence. One key virulence factor is a polysaccharide capsule that expands greatly during infection. To help define capsule synthesis and fungal biology, we provided cells with many different combinations of host-like signals and sampled the cultures over time for transcriptional analysis. The resulting time resolved data set is by far the largest gene expression resource ever produced for *C. neoformans* (881 RNA-seq samples), further enriched by accompanying capsule images and measurements. It revealed surprising findings, including that rich medium suppresses capsule size regardless of other signals. This landmark data resource will be enormously valuable to the research community as it continues to define the relationships between environmental signals and cryptococcal gene expression, biology, and virulence.

---

Late in the 19^th^ century, several scientific articles described a budding yeast with a distinctive capsule, now called *Cryptococcus neoformans*. Today, we know *C. neoformans* as the main causative agent of a deadly meningitis that kills close to 150,000 people each year worldwide^1^. We also know much about the capsule that surrounds this pathogen, including the chemical structure of the polysaccharides that compose it, its key role in disease, and the fact that it is exquisitely sensitive to environmental conditions.^2^ Upon entry to a mammalian host, the capsule dramatically increases in thickness, from a barely perceptible structure to a distinctive shell whose thickness can exceed the cell’s diameter.^3^ Enlarged capsules inhibit phagocytosis of the yeast by host immune cells and shed capsule polysaccharides inhibit host defenses.^4^ The importance of this material in cryptococcal pathogenesis is amply supported by the reduced virulence of strains in which capsule is altered or dysregulated.^2^

Multiple *in vitro* conditions induce the growth of cryptococcal capsule.^5-10^ Conditions that reflect aspects of the mammalian host environment are of particular interest, such as those that incorporate tissue culture medium (TCM) or mammalian serum^9^, human body temperature (37 °C), physiological pH (7.35-7.45),^6,9-12^ host-like CO_2_ concentrations (∼5%)^6^, or nutrient limitations typical of the host environment.^7^

Cyclic AMP (cAMP) signaling is required for capsule growth and virulence of *C. neoformans*.^3,13^ This pathway has been well described in *Saccharomyces cerevisiae*^14,15^ and much of the machinery is also found in *C. neoformans*.^13^ Cellular levels of cAMP reflect its formation from ATP by Cac1 (adenylyl cyclase) and degradation by Pde1 and Pde2 (phosphodiesterases). cAMP binds the repressive subunit of the protein kinase A (PKA) complex, Pkr1, causing it to separate from the catalytic subunit, Pka1. This activates Pka1, allowing it to phosphorylate transcription factors that are central in capsule regulation, including Nrg1^16^ and Rim101^17^. In growth conditions where capsule is normally induced, strains lacking Cac1^3,18^ or Pka1^17,19^ fail to do so. In the same conditions Pkr1-deficient mutants make larger capsules than wild-type (WT) cells^3,20^.

In this paper, we assess key features of capsule inducing conditions, which we call *signals*, focusing on five of them: tissue-culture medium (DMEM or RPMI), temperature (37 °C), CO_2_ (5%), exogenous cAMP, and the addition of buffer. We previously showed that 1,10-phenanthroline, which inhibits transcription, completely blocks capsule growth.^21,22^ We have now systematically explored the effects of these signals on gene expression, cell size, and capsule size, both individually and in combination, over time. We also assessed the effects of deleting several cAMP pathway genes. Our analysis of almost 900 RNA-seq samples and over 5,000 micrographs has allowed us to trace capsule-inducing signals through their effects on gene expression to their ultimate effects on capsule size. This large, uniformly collected dataset, which will be a significant resource for the research community, also enabled us to identify the changes in gene expression that are most consistently associated with capsule growth across a wide range of growth conditions and genetic perturbations. We further discovered that capsule induction is blocked by rich medium under multiple growth conditions and identified new genes that are required for normal capsule thickness.

## RESULTS

### Effects of signals on capsule size

To tease out the effects of various environmental signals on capsule size, we examined all possible combinations of the following growth condition variables: medium (YPD, DMEM, RPMI), CO_2_ (room air, 5%), temperature (30 °C, 37 °C), HEPES buffer pH 7.0 (none, 25 mM), and cAMP (none, 20 mM). For these studies we used *C. neoformans* strain KN99α, which was derived from the reference strain H99.^23^ For rigor of the studies, we grew at least four replicates in each combination of conditions and measured capsule sizes for fifteen fields of view per replicate (totaling 96 annotated cells on average).

We first examined how each individual variable affects capsule size, adjusting for the independent effects of all other variables. To do this, we built a linear regression model with the environmental variables as predictors of capsule size. The resulting regression coefficients showed that the factor with the biggest impact on capsule size was RPMI medium, which increased capsule width an average of 1.12 µm, followed by DMEM (0.67 µm), cAMP (0.53µm) and CO_2_ (0.51 µm), (Fig. 1A).

**Figure 1.**
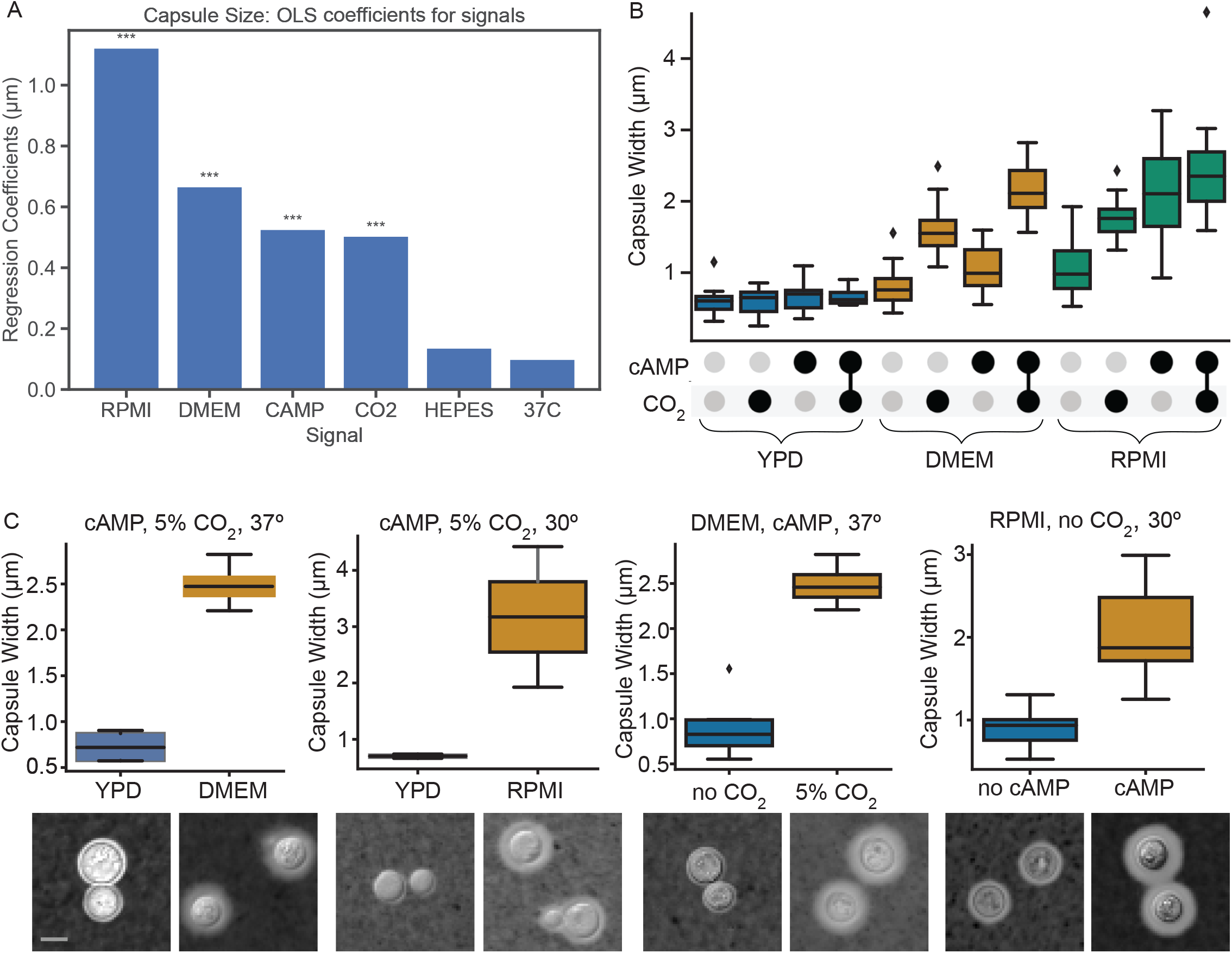
(A) Average effects of environmental signals on capsule size, relative to YPD, in a linear model without interaction terms (N = 232). *** P<0.001 compared to YPD. (B) Capsule sizes for all combinations of media, cAMP, and CO_2_ signals. Horizontal line: median; box: quartile 1 to quartile 3; fence: 1.5 times quartile range; diamonds: individual outlier points. (C) Conditions in which individual signals had their biggest effects on capsule size. None of these conditions includes HEPES buffer. India ink-stained micro-graphs of cells grown in each condition, chosen to match the mean capsule width, are shown below. All images are to the same scale; scale bar, 5mm.

We observed no statistically significant effect of increasing temperature from 30 °C to 37 °C or adding HEPES buffer (Fig. 1A). We therefore combined replicates, regardless of temperature or HEPES, and plotted capsule size for all possible combinations of medium, CO_2_, and cAMP (Fig. 1B). Strikingly, there was no capsule induction in any condition with YPD, regardless of CO_2_ or cAMP addition. DMEM alone yielded negligible induction, but adding cAMP, CO_2_, or both yielded progressively thicker capsules. Results in RPMI were similar, except that it produced larger capsules than DMEM in every condition. In RPMI, cAMP also had a larger effect than in DMEM, such that its effect was similar to that of CO_2_.

To further explore how the signals that affect capsule size interact, we identified the conditions under which each signal had the greatest impact. DMEM and RPMI each had their biggest effects in the presence of both CO_2_ and cAMP. cAMP had its biggest effect in RPMI without CO_2_ at 30°, while CO_2_ had its biggest effect in DMEM with cAMP at 37 °C (Fig. 1C). These observations guided our choice of media in the analyses below.

Taking these results together, the most surprising result was that no combination of temperature, CO_2_, cAMP, or HEPES buffer induced capsule at all in YPD (Fig. 1B; all P-values > 0.2). This suggested to us the novel idea that this rich medium has a repressive effect on capsule (see Discussion). Our data set also allowed us to make several other observations. First, RPMI generally led to greater capsule width than DMEM, holding all other variables constant (linear model, P<10^−8^). Second, CO_2_ or cAMP each increased capsule size in either tissue culture medium (DMEM ± CO_2_, P<10^−3^; RPMI ± CO_2_, P<10^−3^; DMEM ± cAMP, P<0.058; RPMI ± cAMP, P<10^−3^), and the two together yielded the largest capsules.

### Effects of signals on gene expression

To examine the transcriptional response to various combinations of signals, we collected samples for RNA-Seq upon initial exposure to the signals and after 30, 90, 180, and 1440 min. After quality control (see Methods), we were left with RNA-Seq data for 720 samples. These included at least four replicates for each combination of environmental signals, which had been grown and libraries prepared on separate days to account for day-to-day variability in both processes. Each growth batch and library preparation batch also included a control culture grown in our standard non-inducing laboratory conditions: YPD, 30°, no CO_2_, no cAMP, no added buffer. This design allowed us to regress out batch (date) effects, which often dominate experimental factors in large-scale gene expression studies.

To assess the magnitude of signal effects on overall gene expression, we computed principal components and plotted PC1 and PC2 for each sample. Time of growth had the biggest effect on both PC1 and PC2 (Fig. 2A), as can be seen by the progression towards the upper right of data points from successive sampling times. The next greatest effect was due to medium, as shown for the 24 h timepoints in Fig. 2B, where YPD samples (blue) are clearly separated from those in TCM (green and orange). Samples grown in TCM with various combinations of cAMP or CO_2_ were also distinct in terms of transcriptional response (Fig. 2C, showing PC1 and PC3). Together, these results show that the primary differences in gene expression are driven by experimental factors, rather than batch effects, demonstrating the quality of the data. Furthermore, TCM, which has the greatest effect on capsule size at 24 h, also has the greatest effect on gene expression at 24 h.

**Figure 2.**
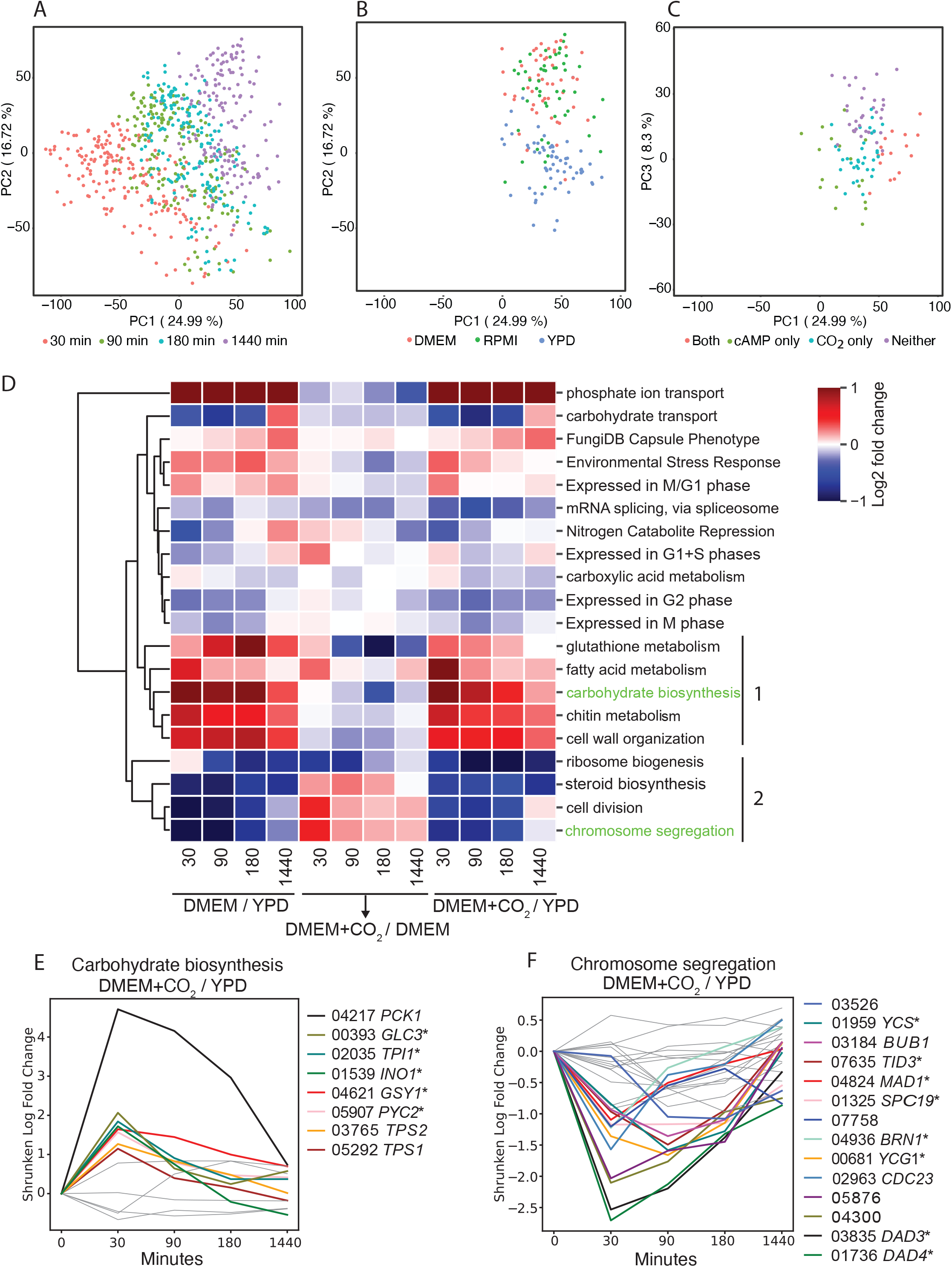
(A-C) Principal Components (PC) plots. (A) PC1 and PC2. (B) 24 h samples only, PC1 and PC2. (C) 24 h samples in tissue culture medium (TCM) only, PC1 and PC3. (D) Log_2_ fold changes (LFCs) for all genes in the indicated categories in DMEM relative to YPD at the same time point (DMEM / YPD), in DMEM with CO_2_ relative to DMEM without CO_2_ (DMEM + CO_2_ / DMEM), or in DMEM with CO_2_ relative to YPD (DMEM + CO_2_ / YPD). Colors indicate the average LFC for all genes in the category, including those that are not significantly differentially expressed. Most LFCs were in the −1 to +1 range; in order to maximize visual discriminability those outside that range were truncated at −1 or +1. Lower case: GO terms; Capitalized terms: other gene sets (see Supplemental File S5 and Methods). Group 1: processes that are strongly upregulated by DMEM; Group 2: processes that are strongly downregulated by DMEM. Green text highlights categories that are expanded in subsequent panels. N=695 samples (E-F) LFCs of individual genes (denoted by the numeric part of CKF44 gene IDs) that are annotated as being involved in carbohydrate synthesis (C) or chromosome segregation (D). Colored lines indicate genes with absolute LFC > 1 at some time point. Asterisk: Gene name provided is that of the *S. cerevisiae* ortholog. N=695.

### Effects of DMEM and CO_2_ on gene expression

In the analyses above, capsule growth required both a tissue culture medium and either CO_2_ or cAMP. To identify the biological functions of the genes that were most responsive to these signals, we started with the response to CO_2_ in DMEM, a combination that had large effects on capsule width (Fig. 1). To assess transcriptional changes we ran DESeq2^24^ on these datasets with a linear model that predicts log normalized gene expression levels from signals and extracted shrunken log fold changes in response to each signal (see Methods).

Next, we carried out over-representation analysis on the most responsive genes at each time point using Gene Ontology (GO) biological process terms together with additional functional categories of interest. Examples of enriched biological functions are shown in Figure 2D, as a heatmap showing the average expression levels of all genes annotated with each term (not just the significantly differentially expressed genes). For each time point, we show the effects of DMEM relative to YPD, the effects of DMEM+CO_2_ relative to DMEM alone, and the combined effects of DMEM+CO_2_ relative to YPD alone.

We first examined genes that are upregulated by DMEM alone. The genes that were most strongly induced are involved in phosphate transport (Fig. 2D, top row), consistent with recent reports that these genes are induced by another capsule-inducing growth condition, 10% Sabouraud’s dextrose medium.^25^ Other biological functions that were strongly upregulated in DMEM include environmental stress response (Fig. 2D, fourth row) and the functions designated as Group 1 (Fig. 2D; see gene lists in Supplemental File S5). This group includes several functions that are notable for their involvement in cell wall synthesis (cell wall organization and chitin metabolism) as well as carbohydrate biosynthesis. The individual expression patterns of each gene in this category are shown in Fig. 2E, plotted as the ratio of expression in the indicated conditions at each time point (so they may reflect differences in either or both conditions). The expression ratio for many of these genes was upregulated by at least 2-fold (colored lines), including for those involved in gluconeogenesis (*PCK1, TPI1, PYC2*), glycogen synthesis (*GLC3, GSY1*), and trehalose synthesis (*TSP1* and *TSP2*, both of which are required for virulence in *C. neoformans^26^*). All of these genes showed peak expression difference from YPD at 30 minutes with a slow convergence back to YPD levels over time (see Discussion).

Another category of interest was genes implicated in capsule synthesis (genes annotated with a capsule phenotype in FungiDB, shown in the third row of Fig. 2D). These were, on average, upregulated by DMEM and barely affected by CO_2_ (see Discussion). Finally, we noted that homologs of yeast genes expressed specifically during the M/G phases of the cell cycle (identified in ref.^27^) were upregulated by DMEM and slightly down regulated by CO_2_, while those expressed at other cell cycle phases were generally downregulated by DMEM, relative to YPD (see Discussion).

Overall, the gene sets that were most strongly down regulated in response to DMEM (Fig. 2D, group 2) are growth related: ribosome biogenesis, steroid biosynthesis, cell division, and chromosome segregation. Expression of each individual gene in the last category (Fig. 2F) shows that, like upregulated genes, the most dramatic differences from YPD levels occurred by the first time point (30 min), with expression in the two cultures becoming more similar by the end of the 24-hour period. The category of carbohydrate transport was also strongly down-regulated relative to YPD for most time points, but then up-regulated at 24 hours (Fig. 2D, second row). Among these carbohydrate transport genes are *LPI8*, whose product promotes cryptococcal uptake by phagocytes^28^, and *GMT2*, which encodes a GDP-mannose transporter required for the synthesis of capsule and other glycoconjugates.^29^

Interestingly, the effect of CO_2_, when added to DMEM, was in the opposite direction from the effect of DMEM relative to YPD for most gene sets in the heat map. However, the CO_2_ response was weaker, so the overall effect of DMEM+CO_2_ was in the same direction as that of DMEM alone. Exceptions to this pattern include ribosome biogenesis and mRNA splicing, both of which were consistently reduced.

### Effects of cyclic AMP on capsule and cell size

Cyclic AMP signaling is essential for capsule growth.^3,13,18^ To investigate the effects of exogenous cAMP on capsule size and gene expression, we performed experiments in conditions in which capsule is normally small, but addition of cAMP generates much larger capsules: RPMI, 30°, room air, no HEPES (Fig. 1B). We found that increasing cAMP yielded a consistent dose response in capsule width (Fig. 3A), with the logarithm of exogenous cAMP concentration a highly significant predictor of average capsule thickness (P<10^−10^). On average, increasing cAMP by a factor of 1.8 increased the capsule width by 0.2 um, although the final step from 11 mM to 20 mM had a much bigger effect (Figure 3A).

**Figure 3.**
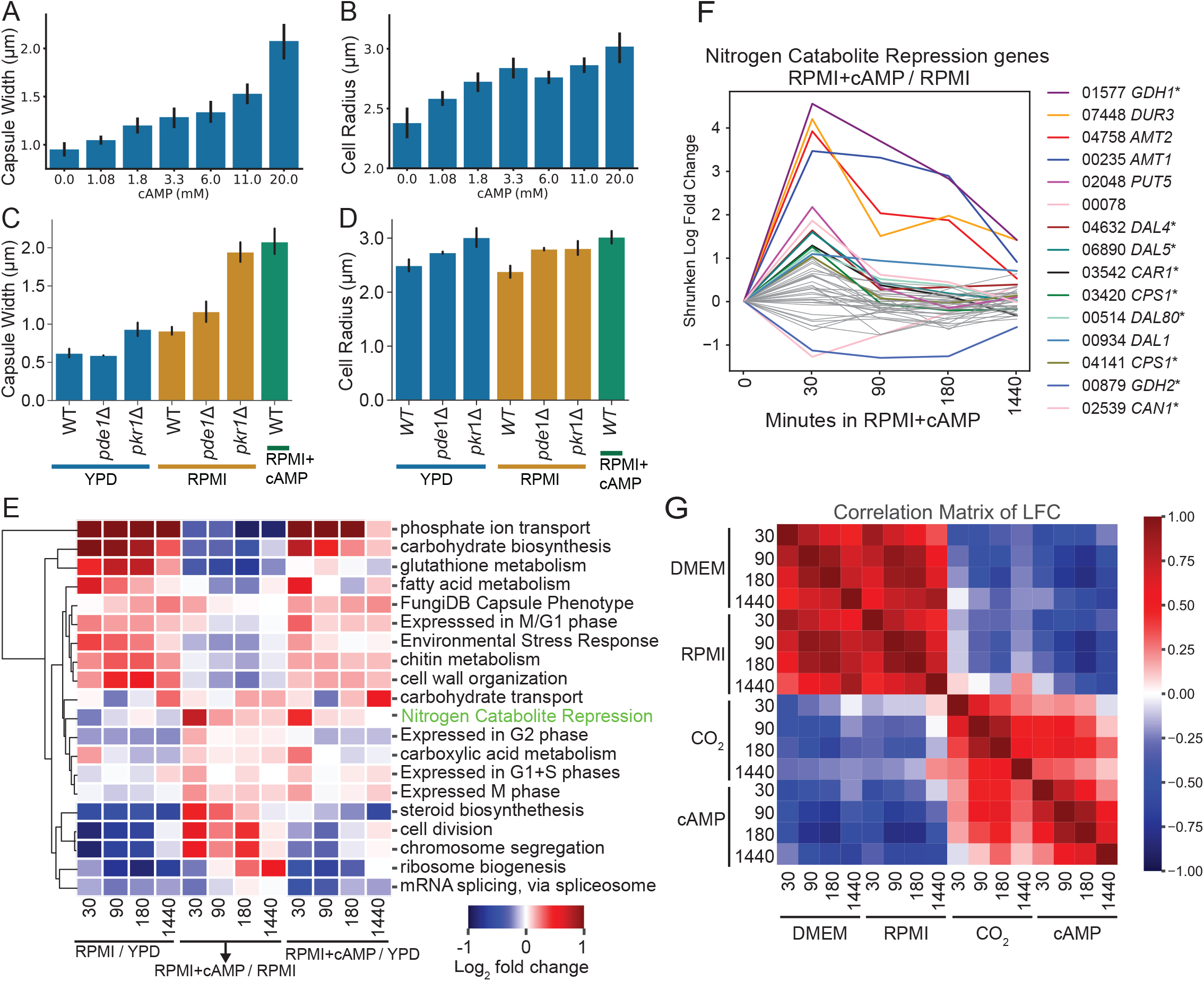
(A) Capsule width or (B) cell body radius as a function of added cAMP. Mean −/+ SD is shown. (C) Capsule width or (D) cell body radius in the indicated strains. Each bar represents 2-3 independent experiments done on different days. (E) Log_2_ fold changes (LFC) for all genes in the indicated functional categories in RPMI (no cAMP) relative to YPD at the same time point (RPMI / YPD), in RPMI with 20 mM cAMP relative to RPMI without cAMP at the same time point (RPMI + cAMP / RPMI), or in RPMI with 20 mM cAMP relative to YPD at the same time point (RPMI + cAMP / YPD). Colors indicate the average LFC for all genes in the category, including those that are not significantly differentially expressed. LFCs were truncated at −1 and +1 to maximize visual discriminability in that range. (F) LFCs of individual genes (denoted as in Fig. 2) that are repressed by the Nitrogen Catabolite Repression (NCR) pathway. Colored lines indicate genes with absolute LFC > 1 at some time point. (G) Correlations between the effects of tissue culture media, CO_2_, and cAMP on expression of all genes. DMEM and RPMI are relative to YPD, all in room air. CO_2_ and cAMP are added to DMEM and RPMI, respectively, and compared to the tissue culture medium alone. Panels E-G are based on a model with 695 RNA-Seq samples.

Exogenous cAMP also increased the average cell body radius, excluding the capsule, (Fig. 3B; ANOVA P < 3*10^−5^), with a 1.8 fold increase in cAMP causing an average increase of 0.11 um (Fig. 3B). Unlike the effect on capsule width, this effect was greater at lower concentrations. Notably, the effect of cAMP on cell size does not explain its effect on capsule width, since the capsule index (capsule width as a fraction of total radius) increased with cAMP concentration (Fig. S1).

We took advantage of mutants in the cAMP pathway to further investigate the role of cAMP in capsule enlargement. We first measured the capsules of cells lacking *PDE1*, which encodes a phosphodiesterase capable of degrading intracellular cAMP. We observed little effect on capsule width in either YPD or RPMI (both at 30° with no CO_2_ or cAMP; Fig 3C), consistent with previous reports^30^. We next examined cells lacking *PKR1*, which encodes the repressive moiety of the Protein Kinase A (PKA) complex. cAMP causes Pkr1 to dissociate from the complex, activating the kinase moiety, so a *pkr1* deletion mutant might be expected to have an enlarged capsule. Indeed, we saw that, when grown in RPMI alone, the *pkr1* mutant’s capsule was similar to that of WT grown in RPMI with 20 mM cAMP (Fig. 3C). We also saw an increase in capsule width of *pkr1* grown in YPD. No other perturbation we tested, including 20 mM exogenous cAMP, yielded increased capsule size in YPD (Fig. 1B; see Discussion). Meanwhile, cell body size was only modestly changed in the mutants in each growth condition (Fig. 3D).

### Effects of cAMP on gene expression

In our experiments, cAMP had the greatest impact on WT capsule size in RPMI with no CO_2_ (Figure 1C, right-most panel). For this reason, we characterized gene expression changes over time in response to these conditions (Fig. 3E), examining the same functional categories we had for DMEM and CO_2_ (Fig. 2D). The responses to RPMI and DMEM were broadly similar (compare Figs. 3E and 2D). Furthermore, adding cAMP or CO_2_ tended to moderate the responses to TCM more often than it reinforced them. However, we did observe some differences. For example, while RPMI and DMEM both reduced the expression of ribosome biogenesis genes, adding CO_2_ reinforced that effect at all time points, while adding cAMP moderated it at later time points. We also noted that genes involved in carboxylic acid metabolic processes (mainly nitrogen assimilation and amino acid metabolism) responded differently to RMPI+cAMP and DMEM+CO_2_, especially at early time points.

Nitrogen source is another factor that has been implicated in capsule regulation. ^6^ Nitrogen catabolite repression (NCR) is a process that represses the expression of certain genes when preferred nitrogen sources, such as glutamine (the main nitrogen source in DMEM and RPMI) are available.^31^ In RPMI, the genes in this category were indeed slightly repressed compared to YPD (Fig. 3E, left columns). However, adding cAMP released NCR rapidly, most notably at 30 min (Fig. 3E, middle columns), suggesting that the cells responded to cAMP as though they had been moved to a less preferred nitrogen source (despite the presence of glutamine in the medium; see Discussion). This is consistent with the general role of cAMP in cryptococcal stress response. cAMP addition also overcame the RPMI effect, so that their combined effect was to relieve NCR (Fig. 3E, right columns). The detailed effect of adding cAMP to RPMI is shown in Fig. 3F.

Finally, to evaluate the effects of signals across all genes, rather than on specific processes, we calculated the global correlations between pairs of signals at each time point. As we had observed for selected groups of genes (Fig. 3E), the effects on gene expression of the two tissue culture media (DMEM and RPMI) are highly correlated (Fig. 3G). The effects of CO_2_ and cAMP were positively correlated with each other, consistent with previous evidence that CO_2_/HCO_3_ stimulate adenylyl cyclase, which generates cAMP.^32^

### Expression of multiple genes is associated with capsule size

Next, we set out to identify genes whose expression is strongly associated with capsule development. Our approach was to dichotomize capsule size into “induced” or “not induced” and the expression of each gene into “high” or “low” and analyze the relationship between expression and capsule induction for each gene separately. We considered a sample to be “induced” if its median capsule width after 24 hours exceeded 1.13 um, three standard deviations above the mean of all WT cells in YPD. For each gene at each RNA-Seq time point, we then searched for a threshold to divide high and low expression for that gene that was most associated with induction status. Specifically, we calculated the *X*^2^ statistic for association between induction status and gene expression status (Fig. 4A; see Methods for details). We were not concerned with P-values here, but with using the statistic itself to rank genes by their association with capsule size. The maximum *X*^2^ value, over all time points, was assigned to the gene and the genes with high *X*^2^ values were identified as potentially involved in capsule induction (Table S1). However, we were mindful that these associations can arise in many ways and do not necessarily reflect a causal role in capsule induction.

**Figure 4.**
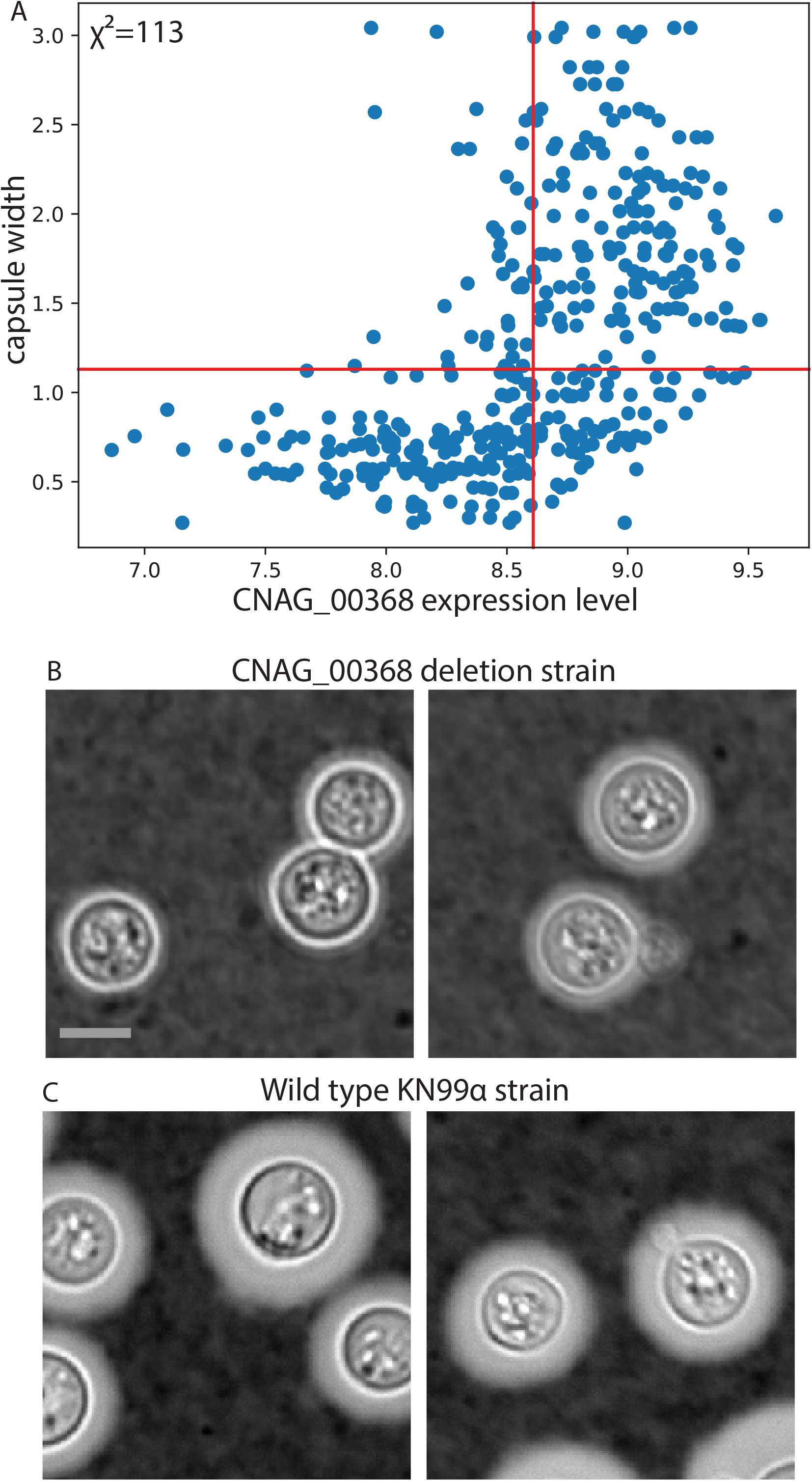
(A) Example gene with high *χ*^2^, reflecting the visual relationship between its expression level and capsule size. Cells with capsules larger than the horizontal lines are considered induced while genes with expression levels to the right of the vertical red lines are considered highly expressed. Expression levels are log_2_ of counts, +1 pseudo count, after normalization by DESeq2 and batch effect removal. (B-C) Example micro-graphs of cells of the indicated strain, stained with India ink. All images are to the same scale; scale bar, 5 mm.

### New genes whose deletion affects capsule size

As noted above, the same conditions that induce capsule and stimulate changes in expression of genes essential for capsule growth also affect expression of genes involved in multiple other processes, including cell division, nutrient utilization, or other cellular processes (see Figs. 2D and 3E). We therefore expected only a small subset of the genes whose expression is correlated with capsule thickness to be essential for capsule development. To seek such genes, we examined deletion strains. Mutants corresponding to 24 of the 33 genes in Table S1 were available in a publicly available deletion collection^33^ or in our own collections. We grew each mutant in the conditions we found to induce large capsules (RPMI, 37 °C, 5% CO_2_, no cAMP) and measured their capsules using our rigorous protocols (see Methods).

We were excited to identify a mutant that was not previously known to influence capsule width. Deletion of CNAG_00368, which encodes a homolog of *S. cerevisiae* Vps53, reduced mean capsule width by 0.44 µm (Fig. 4B,C). A scatter plot of capsule width versus expression for this gene is shown in Fig. 4A.

An adventitious finding in these studies was that a strain deleted for CNAG_05977 had mean capsule width 0.54 um smaller than wild-type cells (see Discussion). This strain was not chosen as described above – we had intended to test CNAG_06050, but routine genome sequencing for quality control showed that the strain we tested in fact had a deletion of CNAG_05977. We verified the genotypes of both deletion strains by whole-genome sequencing.

## DISCUSSION

This work makes significant contributions in three areas. First, we have generated a major dataset that will be a tremendous resource for researchers in the field. Second, we have used these data to make interesting observations about how cells react to environmental conditions that induce capsule. Third, we identified a set of genes whose expression correlates with capsule size and discovered two new genes that influence capsule thickness.

When we undertook this project, it was known that capsule growth requires both new transcription^6,22^ and an intact cAMP/PKA pathway.^3,13,18^ It was also known that, in tissue culture medium (TCM), capsule can be induced by increasing the concentration of dissolved CO_2_/HCO_3_, achieved by growth in a high CO_2_ atmosphere, addition of NaHCO_3_, or both.^6^ We set out to determine how various combinations of capsule inducing signals would affect gene expression and capsule size. This led to a massive, freely available dataset consisting of RNA-Seq in biological quadruplicate in 42 combinations of potentially capsule-inducing signals, as well as cAMP titrations and mutants in the cAMP signaling pathway (881 total RNA-Seq samples). Each RNA-Seq experiment comes with matched India ink images and capsule width measurements. The dataset also includes images and capsule-width measurements for selected gene deletion mutants, yielding a total of 47,458 annotated cells in 5,175 images of 392 biological samples.

We found that either a 5% CO_2_ atmosphere or cAMP could induce capsule in TCM, but YPD, a rich medium containing yeast extract and peptone, completely blocks capsule growth. This repressive effect is upstream of PKA activation, since deleting *PKR1* increases capsule sizes of cells growing in YPD (Fig. 3C). However, adding 20 mM exogenous cAMP, which is thought to act directly on the Pka1/Pkr1 complex, does not enlarge the capsules of cells growing in YPD (Fig. 1B). It may be that cAMP cannot completely inactivate Pkr1 or that cells do not maintain a high enough internal concentration for long enough to fully inactivate it.

RPMI alone caused capsules to enlarge slightly, whereas DMEM alone produces a barely detectable enlargement (Fig. 1B). RPMI contains less of most nutrients than DMEM, making it a generally more stressful condition. One possible explanation for these observations is the absence of iron in RPMI (DMEM contains 10^−4^ g/L ferric nitrate). These results are reminiscent of observations concerning titan cells, extremely large cells that form during infection and specific *in vitro* conditions.^34^ Interestingly, titan cell formation can be induced by low-density inoculation into RPMI at 37 °C with 5% CO_2_, but substitution of DMEM for RPMI prevents this. The critical differences in this case were identified as the presence of iron in DMEM and the presence of para-aminobenzoic acid in RPMI.^35^

In addition to the conditions we studied, capsule growth is known to be stimulated by specific stressors, some of which resemble stresses encountered in a mammalian host. For example, growth in mammalian serum without added nutrients induces capsule growth at 30 °C or 37 °C, with or without CO_2_, although this does not occur in Sabouraud dextrose broth, a rich medium like YPD.^9^ Sabouraud medium alone, diluted to 10% normal concentration and buffered to pH 7.4 (termed CAP medium), is reported to induce capsule.^8^ Iron deprivation in LIM medium, which consists of 5 g/L glucose, 5 g/L asparagine, minerals, and 55 mM EDDA at pH 7.4, can also induce capsule.^7^ Thus, rich media consistently block capsule induction while several forms of nutrient deprivation induce it. Nutrient deprivation is likely the normal state of Cryptococcus, whether in a mammalian host or in the environment, so we suggest that it makes more sense to think of rich media as blocking capsule growth than to think of TCM as inducing it.

Low pH (6.1) can also block capsule induction.^6^ High osmolarity appears to reduce capsule thickness, although there has been speculation about whether this is a cellular response or simply physical compression of capsule.^12^ A cellular response is consistent with the observation that deletion of *HOG1*, a key component of the high osmolarity response, increases capsule size in DMEM+CO_2_ induction.^3,36^ Studying the effects of these stimuli on gene expression may provide additional insights about which changes in gene expression are essential for capsule growth.

Turning to gene expression, tissue culture media (TCM) have several interesting effects. First, they decrease the expression of genes associated with growth and increase that of genes associated with stress. Further, gene expression in TCM suggests an accumulation of cells in the M/G1, post-mitotic phase of the cell cycle. Finally, TCM tends to engage nitrogen catabolite repression, probably because of its glutamine content.

Surprisingly, both CO_2_ and cAMP moderate or reduce most of the effects of TCM on gene expression. The fact that either of these components can act as the necessary partner with TCM for capsule induction, but their overall effects on gene expression are opposite that of TCM, poses intriguing questions: When CO_2_ is added to TCM, which of the resulting gene expression changes are critical for capsule induction and do those changes work by dialing down the effects of TCM on key genes, by reinforcing the effects of TCM on the few genes that respond in the same direction, or by acting on genes that do not respond to TCM?

Both phosphate and nitrogen availability have been suspected as regulators of capsule size. Both nutrients activate sensors that stimulate the PKA pathway.^37-39^ It has also been reported that expression of phosphate acquisition genes is strongly induced in CAP medium and that subsequent addition of KH_2_PO_4_ reduces capsule width as well as cell size.^10^ We found that phosphate transporters were also strongly induced by TCM, although this effect was much reduced by addition of CO_2_ or cAMP.

Regarding nitrogen availability, it has also been reported that YNB medium (trace nutrients, (NH_4_)_2_SO_4_, and glucose) with bicarbonate and 5% CO_2_ induces capsule, but only if the (NH_4_)_2_SO_4_ is replaced by arginine.^6^ Both are considered preferred nitrogen sources for *S. cerevisiae*^31^, so the cells may be responding to amino acids, rather than the quality of the nitrogen source. Further linking amino acids and capsule, several genes with expression levels that are associated with capsule size encode proteins that act in transport of amino acids or oligopeptides, such as CNAG_01119 and CNAG_02539, which are orthologs of the *S. cerevisiae* oligopeptide transporter *PTR2* and amino acid transporter *DIP5*, respectively (Tables S1).

As mentioned earlier, conditions that induce capsule also impact multiple cellular response pathways. Nonetheless, we hypothesized that we might be able to identify genes required for normal capsule size among those genes whose expression was best correlated with capsule thickness and tested this using deletion mutants. Unfortunately, there is no deletion mutant in the Madhani collection for many of the genes whose expression is most predictive of capsule thickness (Tables S1). Our own attempts to delete five of these genes also failed, suggesting that they may be essential. Studying their effects on capsule thickness, therefore, may require construction of over-expression strains or development of more robust tunable promoter systems than are currently available for *C. neoformans*.

When a mutant was available for a gene whose expression was statistically associated with capsule thickness, we assayed it for this characteristic. Unsurprisingly, most mutants did not show a significant change in capsule size. Some of the corresponding genes may indeed have no role in capsule elaboration, their correlation with capsule size resulting from parallel pathways activated by the same stressful conditions that lead to capsule growth. Others may play a role in capsule growth that can also be filled by other proteins. Deletion of still others may affect capsule in ways that our methods do not detect, such as changes in glycan composition, structure, or other features. An alternative approach that hypothesizes capsule phenotypes for genes whose expression pattern is similar to those of known capsule genes was recently reported to be effective, with 6 of 12 tested mutants having altered capsule width.^40^

We did identify two genes, not previously known to influence capsule, whose deletion affects capsule size in inducing conditions. Deletion of CNAG 00368 reduced mean capsule thickness by 0.44 um. This gene encodes an ortholog of *S. cerevisiae* Vps53, which is involved in recycling proteins from endosomes to the late Golgi. Vps53 not been studied in Cryptococcus, but a protein that is part of a different complex involved in recycling proteins from endosomes, Vps23, is known to be required for capsule elaboration.^41^ Reinforcing the theme of protein and peptide recycling, deletion of CNAG_05977 (proteasome activator subunit 4) reduced capsule size by 0.54 um. Future work will be needed to define the specific mechanisms by which these genes influence capsule.

Our dataset of matched RNA-Seq time courses and capsule thickness measurements has yielded insights into the environmental signals and gene expression changes that affect capsule size. However, this is only the beginning. We expect that future analyses by our group and other researchers will yield additional insights, not only into capsule biology but into numerous aspects of cryptococcal physiology, many with importance for disease.

## METHODS

Additional details for all methods can be found in the online supplement.

### Cell growth

To maximize reproducibility of RNA-seq and capsule imaging studies, cell recovery from frozen stocks, initial culture in YPD, inoculation into preconditioned media, and growth followed strictly controlled protocols. These methods are detailed in the Supplemental Material.

### Microscopy and manual image annotation

1-ml samples were collected for imaging from the stock cell suspension prior to inoculation of flasks or from cultures at 1440 min, fixed, resuspended in PBS, and mixed with India ink (5 parts cells:2 parts ink) for brightfield microscopy. Images were manually annotated using a custom annotation interface written in Mathematica / Wolfram Language (available on request). Fifteen fields were annotated for each replicate of each combination, yielding an average of 96 annotated cells per replicate (min 31; max 361; standard deviation 54).

### Analysis of capsule thickness in gene deletion mutants

Effects of deletion mutants on capsule thickness were evaluated using a Linear Mixed Models framework with individual cells’ thicknesses as the datapoints. The model had fixed effects for intercept and genotype; each specific biological sample was used as a grouping variable with a random intercept for each group.

### RNA-Seq

Libraries were constructed using the NEBNext Ultra Directional RNA Library Prep Kit from Illumina and samples were pooled at 10 nM. The pools were sequenced on a NextSeq 500 using the High75v2 kit as 1×75 (single end).

### RNA-Seq Computational Pipeline and Analysis

Documented code implementing the process described below is available on our public github repository: https://github.com/BrentLab/brentlabRnaSeqTools. Reads were aligned with Novoalign (version 4.03.02) and quantified with HTSeq (0.9.1) using the FungiDB KN99α genome sequence (FungiDB accession ASM221672v1). Custom scripts were used to verify the strain in each sample by calculating coverage over the open reading frame of the putatively deleted genes and over marker genes, ensuring that the former were absent and the latter present. Documented code and parameters for strain validation and QC can be found here: https://brentlab.github.io/brentlabRnaSeqTools/articles/QC_Library_Quality.html#crypto coccus. RNA-Seq samples that passed strain validation were subjected to two phases of QC. In Phase 1, files were labeled as ‘passing’ if they contained at least 10^6^ reads aligned to protein coding regions and less than 7% of all reads failed to align.

In Phase 2, we evaluated replicate agreement using the Regularized Log Expression (RLE) ^42^. First, we used DESeq2 (version 1.34.0) to estimate the effect of the library date (the known batch effect). We removed the batch effect using the DESeq coefficients for the library dates such that the data were standardized to a single date, resulting in adjusted expression levels on a log_2_ normalized scale. To compute the RLE value for a given gene, the median expression level of that gene, across all samples in a replicate set, was subtracted from the expression level of the gene in each sample. For each sample, we then calculated the interquartile range of the distribution of these RLE values across genes. If the interquartile range of RLEs of a given sample was greater than 1, indicating that more than half of genes deviated from their respective medians by a factor of 2 in adjusted count, the sample was considered an outlier and failed for replicate agreement. 22 samples (2.75%) failed replicate agreement and were discarded.

### Gene set overrepresentation analysis

Differential expression analysis was conducted using DESeq2 with a model containing the main effect of time, represented as a categorical variable, and the interaction effects of time with environmental signals. The interaction effects indicate how the normal effect of time in standard laboratory conditions (YPD, no CO_2_, 30 °C, no cAMP, no HEPES) is modulated by the presence of other factors at each time point. For each time-point:signal interaction, a gene was considered to be differentially expressed if the absolute value of its shrunken log_2_ fold change (LFC) reached 1.0. (DESeq shrinkage reduces the estimated LFCs of factors with greater replicate-to-replicate variance). Overrepresentation analysis was carried out using GOTermFinder (see online supplement for additional details).^43^ Gene annotations came from UniProtKB, release 2020_01, and were supplemented with several annotations derived from literature (see Online Supplement and File S5).

### Selection of potential capsule-associated genes: Chi-squared method

The following procedure was carried out separately for samples at each time point. To calculate the *X*^2^ statistic for each gene, we first classified all samples with average capsule width greater than 1.13 μm (3 S.D. above the mean of all YPD conditions) as induced. Other samples were classified as uninduced. Samples were also classified by high or low expression of each gene, yielding a 2×2 contingency table. For each gene, we tried all possible thresholds of high versus low expression, constructed the contingency table, and calculated the *X*^2^ statistic. Finally, we chose the threshold that yielded largest *X*^2^ statistic. These statistics were only used to characterize the degree of association between the expression of a gene in a sample and that sample’s induction status. They were not used for hypothesis testing.

Genes were ranked for likelihood of the corresponding deletion strain exhibiting a capsule phenotype by the maximum of their *X*^2^ values, across all time points. Genes with high *X*^2^ statistics were selected for testing. For genes that were positively correlated with capsule size, the gene deletion mutant was tested in inducing conditions (RPMI, 37 °C, 5% CO_2_) with the prediction that deletion would reduce capsule size compared to WT grown in the same conditions. For negatively correlated genes, the gene deletion mutant was tested in non-inducing conditions (YPD, 37 °C, 5% CO_2_) and in ‘almost-inducing’ conditions (RPMI, 37 °C, room air); the rationale for the latter was that the conditions were quite close to inducing conditions and perhaps favor capsule synthesis.

### Forming metagenes as features for machine learning

From the original gene expression data matrix, we sought to decrease the number of features by filtering and combining genes. We first removed low variance genes by filtering out genes whose expression in >= 95% samples was within one log_2_ of their mean, indicating that less than 5% of samples showed substantial changes. We next combined highly correlated genes into metagenes (correlation threshold > 0.8). The “expression level” of a metagene was then set to the mean of the expression levels of its constituent genes. We repeated this process to combine genes into metagenes until we had combined all highly correlated genes into metagenes.

## AVAILABILITY OF DATA AND MATERIALS

All RNA-Seq data are available from NCBI GEO under accession numbers GSE226255, GSE226637, or GSE226651, which are subseries of GSE226656. All capsule size measurements are available as supplemental files. Corresponding India ink images and mutant strains are available on request.

