## Supplementary material for "Leveraging a new data resource to define the response of *C. neoformans* to environmental signals": Online supplement

**Data availability and access**

All RNA-Seq data (reads and the count tables we used for analysis) are available from NCBI GEO under accession numbers GSE226255, GSE226637, or GSE226651.

These are subseries of GSE226656. To review GEO accession GSE226656, go to <https://www.ncbi.nlm.nih.gov/geo/query/acc.cgi?acc=GSE226656> .

**Supplemental Data files**

Supplemental files S1-S4 file contain capsule and cell size measurements for individual annotated cells and descriptions of genotypes and growth conditions. The four files are:

- File S1 capsule sizes 42 Combinations of Growth Conditions.csv
- File S2 capsule sizes cAMP Titration WT.csv
- File S3 capsule sizes PKR1 PDE1 deletions.csv
- Files S4 capsule sizes of deletion mutants.csv

“File S5 Supplemental Gene Categories” contains the gene sets that were added to the Gene Ontology Biological Process terms when doing over-representation analysis and making heatmaps.

“File S6 expression of capsule implicated genes” contains expression time courses of all genes that were listed in FungiDB as having altered capsule phenotypes. Specifically, their expression as function of time in DMEM+CO_2_ relative to their expression in YPD without CO_2_ at the same time points.

**Supplemental Figures**

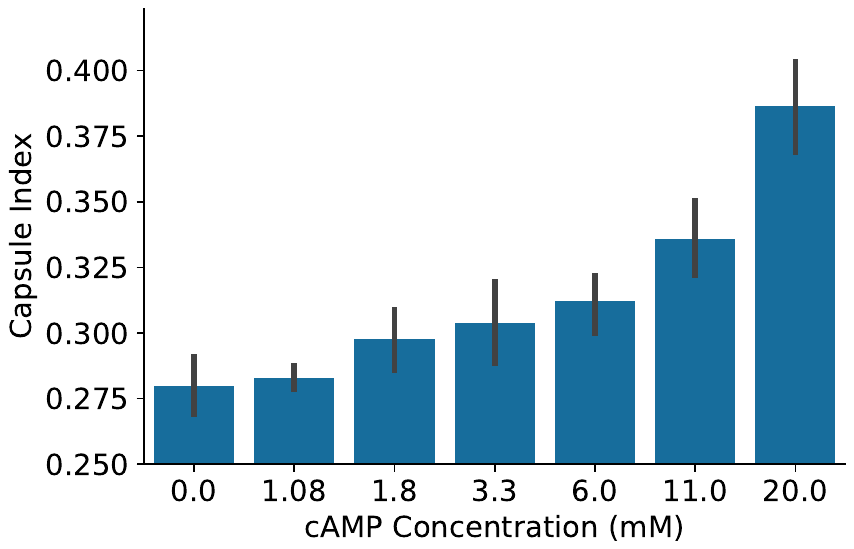

**Fig. S1.** Same as main text Figure 3A, except that capsule index is shown instead of capsule width. Capsule index is capsule width divided by (cell radius + capsule width), i.e. capsule width as a fraction of total radius.

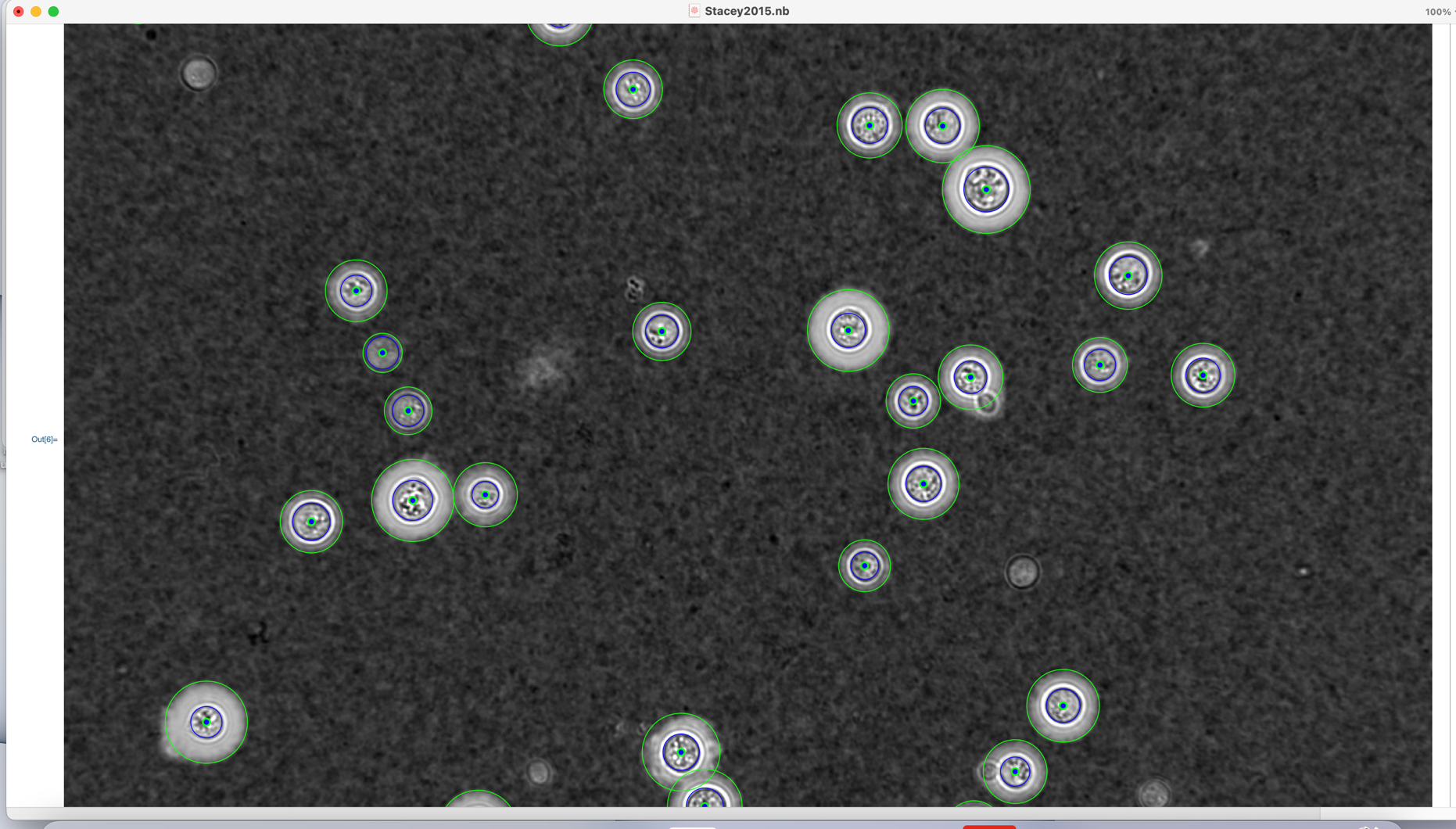

**Fig. S2** Our custom interface for annotating cell and capsule sizes. The interface is written in the Wolfram language (a.k.a. Mathematica). Users must click 3 times to define a perfect circle on the outer edge of the capsule and 3 more times to define a perfect circle on the cell boundary – specifically, we annotate the dark ring between the white wall and the gray cytoplasm. The cell center, cell diameter, and capsule diameter are automatically recorded in a file for later analysis. Annotations of the same fields by different annotators yield highly reproducible capsule and cell sizes.

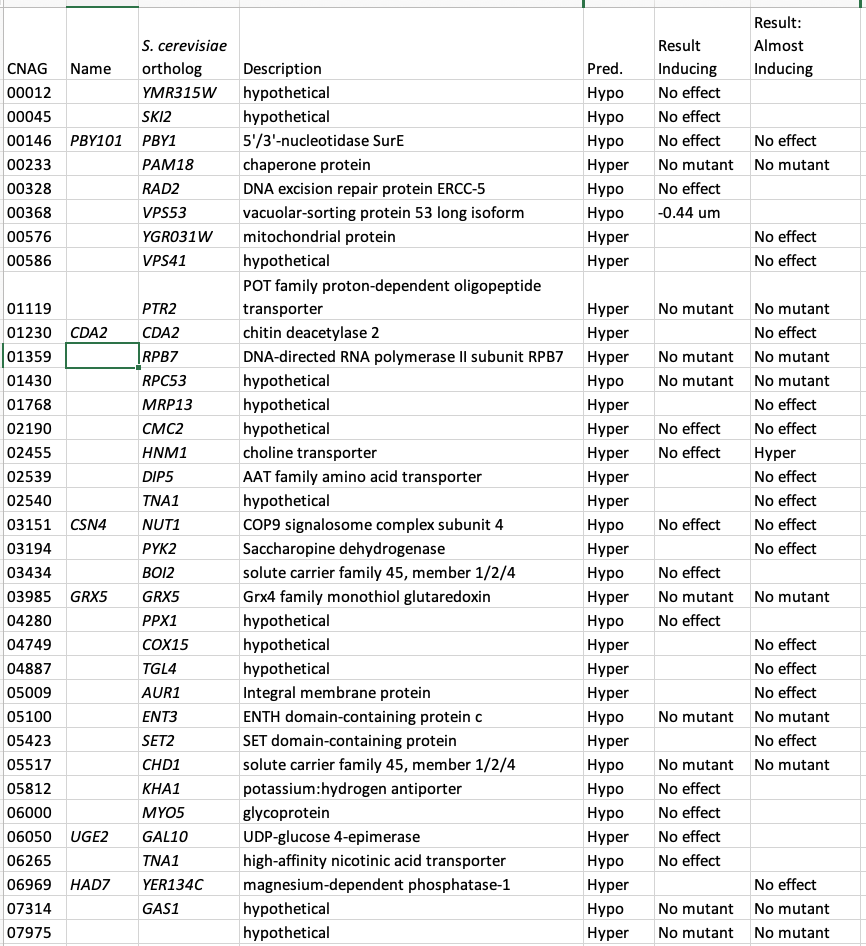

**Table S1** Genes the chi-squared method predicts to be involved in capsule development. CNAG, numerical portion of gene identifier; Pred., predicted capsule effect; Result Inducing; result of growth in inducing conditions; Result: Almost, result of growth in ‘almost inducing’ conditions defined in the main text.

**Supplemental Methods**

Cell growth

Strains were removed from -80 °C to YPD agar, grown at 30 °C for 2 days, and a single colony was inoculated into 4 ml yeast peptone dextrose (YPD) and grown overnight at 30 °C shaking at 230 rpm. One day before the experiment, 1 ml of the ON culture was inoculated into 100 mL YPD and grown as above. At the beginning of the experiment, this culture was spun down, washed in 25 ml room temperature (RT) PBS, and resuspended in 10 ml PBS. 2 x 10^8^ cells were then inoculated into each of four T75 flasks (MIDSCI TP90076) containing 20 ml preconditioned medium (described below) and incubated under the conditions specified for the experiment.

A single flask was harvested for each time point (30, 90, 180, and 1440 min after inoculation) by transferring 18.5 mL of culture to a tube on ice containing 2 ml RNA Stop solution (5% tris-saturated phenol in ethanol). The cells were then sedimented (3000g, 5 min, 4C), the supernatant fraction decanted, and the pellet frozen in liquid nitrogen and stored at -80 °C.

For imaging, 1-ml aliquots of cells were collected prior to inoculation of flasks and at 1440 min, sedimented as above, resuspended in PBS, and counted with a Nexcelom BioScience Cellometer Auto M10. 10^6^ cells were added to 0.5 ml PBS containing 3.7% formaldehyde and the cells were fixed by end-over-end rotation at RT for at least 30 minutes. Fixed cells were stored at 4 °C.

Environmental signals and growth conditions

All flasks were prepared and then preconditioned for 24 hours at the designated temperature before use. Although bicarbonate was not added to the cultures that were exposed to CO_2_, 24 hours is ample time for atmospheric CO_2_ to reach equilibrium with dissolved HCO_3_^--^.^1^ Although this does acidify the medium, the acidification is buffered by HEPES in the HEPES conditions; addition of HEPES buffer also had very little effect on gene expression or capsule size. Cultures were then inoculated as above and incubated without rotation. DMEM (Sigma D6429) and RPMI (Sigma R8758) were used as purchased except in “HEPES” conditions, where sterile 1M HEPES pH 7.0 was added to a final concentration of 25 mM.

### Microscopy and manual image annotation

Fixed cells were sedimented as above and resuspended in 250 μl PBS. 10 μl of this suspension were mixed with 4 μl of India ink and imaged by brightfield microscopy at 63X magnification. Images were manually annotated using a custom interface written in Mathematica / Wolfram Language (available on request). Fifteen fields were annotated for each replicate of each combination, yielding an average of 10^8^ annotated cells per replicate (SD 56).

Cell size censoring

For WT cells across various growth conditions, 21,530 cells were annotated. 4.5% of them had cell radius (excluding capsule) < 1.5 um and 1.7% had cell radius > 4.0 um. Most of these came from a few growth conditions: For small cells, most were from RPMI 30° no CO_2_ no cAMP, YPD at 30° or 37° with no CO_2_ no cAMP, or RPMI 30° 5% CO_2_ no cAMP; for large cells, most came from conditions with cAMP but not CO_2_: RPMI 30, RPMI 37, or YPD 37. Since cell size and capsule size are correlated and our primary interest is the latter, we removed the cells with radii outside the [1.5 um – 4.0 um] interval before further analysis (however, they have not been removed in the datasets provided as a supplementary file). This had no effect on the conclusions. For example, across all deletion mutants tested, censoring did not change the average deletion effect by more than 0.057 μM, far below the level of statistical significance.

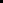

Gene deletion mutants and analysis of capsule thickness

Most gene deletion mutants were from the Madhani deletion collection via the Fungal Genetics Stock Center. For genes of interest where mutants were not present in that or other reported collections, we attempted to delete them at least twice, using our standard split marker biolistic strategy as described in ref.^2^ or CRISPR.^3^ These attempts were considered to have failed if we obtained no colonies or if the colonies obtained, upon analysis, maintained the target gene. Effects of deletion mutants on capsule thickness were evaluated using a Linear Mixed Models framework with individual cells’ thicknesses as the datapoints. The model had fixed effects for intercept and genotype; each specific biological sample was used as a grouping variable with a random intercept for each group.

RNA Isolation

Frozen cell pellets were thawed on ice and 875 μl TRIzol (Ambion 15-596-018) was added. Cells were then lysed by beating with 750 μl 0.5mm silica-zirconia beads for 3 min in a Minibeadbeater (Biospec Products) and RNA was isolated following the manufacturer’s protocol. Residual DNA was removed with the Turbo DNA-free Kit (Life Technologies AM1907) and polyA+ RNA isolated from ~1 μg of total RNA using the NEBNext Poly(A) mRNA Magnetic Isolation Module.

Library preparation and sequencing

Libraries were constructed using the NEBNext Ultra Directional RNA Library Prep Kit from Illumina and samples were pooled at 10 nM. The pools were sequenced on a NextSeq 500 using the High75v2 kit as a 1x75 with 7 bp Index1 and 6 bp Index2 with 1% PhiX. Fastq files were demultiplexed using the Illumina bcl2fastq2 allowing 1 bp mismatch.

RNA-Seq Computational Pipeline and Analysis

Scripts used for RNA-Seq analysis are available at

<https://github.com/BrentLab/brentlabRnaSeqTools>

with instructions and tutorials available here:

https://brentlab.github.io/brentlabRnaSeqTools/index.html.

Reads were demultiplexed into fastq files using Illumina bcl2fastq2 allowing 1 bp mismatch. Resulting fastqs were aligned with novoalign (version 4.03.02) and quantified with HTSeq (0.9.1) using the FungiDB KN99α genome sequence. Genome annotations were augmented with noncoding RNA regions lifted over from the current H99 genome. The quantification for downstream analysis was calculated over gene feature exons, but quantification of the open reading frame (ORF) was also performed for QC purposes. Samtools (version 1.12) was used to convert the SAM output from novoalign to BAM and index Bam files. Novoalign and HTSeq logs were collated by MultiQC (version 1.2).

Custom scripts were used to verify the strain represented in each fastq by calculating ORF coverage over putatively deleted genes and marker genes used to replace them, ensuring that the former were absent and the latter present. Thresholds used in this analysis are documented here:

https://brentlab.github.io/brentlabRnaSeqTools/articles/QC_Library_Quality.html

We found that ORF deletion frequently resulted in increased transcription of UTRs, so full-transcript counts are not appropriate for validating ORF deletions. RNA-Seq samples that passed strain validation were subjected to two phases of QC. In Phase 1, files were labeled as ‘passing’ if they contained at least 10^6^ reads aligned to protein coding regions and less than 7% of all reads failed to align.

In Phase 2, we evaluated replicate agreement using the metric Regularized Log Expression (RLE) ^4^. First, we used DESeq2 (version 1.34.0) to estimate the effect of the library date (the known batch effect). We removed the batch effect using the DESeq coefficients for the library dates such that the data was standardized to a single date, resulting in adjusted expression levels on a log_2_ normalized scale. To compute the RLE value for a given gene, the median log expression level of that gene, across all samples in a replicate set, was subtracted from the log expression level of the gene in each sample. For each sample, we then calculated the interquartile range of the distribution of these RLE values across genes. If the interquartile range of RLEs of a given sample was greater than 1, indicating that more than half of genes deviated from their respective medians by a factor of 2 in adjusted count, the sample was considered an outlier and failed for replicate agreement. 22 samples failed replicate agreement and were discarded.

Gene set overrepresentation analysis

Differential expression analysis was conducted using DESeq2 with a model containing the main effect of time, represented as a categorical variable, and the interaction effects of time with environmental signals. The interaction effects represent how the normal effect of time in standard laboratory conditions is modulated by the presence of other factors (signals) at each time point. For each time-point:signal interaction, a gene was considered to be differentially expressed if the absolute value of its shrunken log_2_ fold change (LFC) reached 1.0. (DESeq shrinkage reduces the estimated LFCs of factors with greater replicate-to-replicate variance in order to make conservative estimates of effects when uncertainty is high.)

Overrepresentation analysis was carried out using GOTermFinder.^5^ Gene annotations came from UniProtKB, release 2020_01. We did not filter annotations by evidence codes. Uniprot was supplemented with several annotations derived from literature (File S2). Literature sources listed genes in *Saccharomyces cerevisiae* that are expressed during specific phases of the cell cycle^6^ or as part of the integrated stress response^7^ or repressed in the presence of preferred nitrogen sources^8^. We identified cryptococcal orthologs of *S. cerevisiae* genes in each of these categories by using the mapping provided in ref.^9^. The corresponding annotations were applied to the *C. neoformans* orthologs producing the gene sets described in Figures 2 and 3 by the terms M/G1, G1+S, G2, M, Environmental Stress Response, and Nitrogen Catabolite Repression. The gene set labeled FungiDB Capsule Phenotype consists of all Cryptococcus genes annotated as having a capsule-related phenotype when deleted. We also added five genes to the set annotated “phosphate ion transport” based on ref.^10^ These additional and augmented gene sets were used for over-representation analysis along with all the gene sets from UniProtKB. Terms with Bonferroni-corrected P-values < 0.05 were investigated for interpretable patterns across signals and timepoints and a subset of terms was selected manually.

### Selection of potential capsule-associated genes: Chi-squared method

The following procedure was carried out separately for samples at each time point. To calculate the 𝜒^2^ statistic for each gene, we first classified all samples from all conditions as either enlarged or unenlarged. To be enlarged, a sample needed a mean capsule width of greater than 1.13 μm, or 3 standard deviations off the mean (across samples) of the means (across cells within a sample) of all samples grown in YPD, regardless of other signals provided. This criterion was chosen because there was a big gap in capsule thickness between all YPD samples and all samples in conditions that we expected to be enlarged. Samples were also classified by high or low expression of each gene, yielding a 2x2 contingency table, in which samples were classified as enlarged or unenlarged and high versus low expressors of the gene. For each contingency table, we calculated the largest 𝜒^2^ statistic. For each gene, we tried all possible thresholds of high versus low expression and chose the threshold that yielded the largest 𝜒^2^ statistic. These statistics were only used to characterize the degree of association between the expression of a gene in a sample and that sample’s induction status. Since the statistics were not used for statistical hypothesis testing, the issue of multiple hypothesis testing does not apply.

Genes were ranked for likelihood of the corresponding deletion strain exhibiting a capsule phenotype by the maximum of their 𝜒^2^ values, across all time points. All genes with max 𝜒^2^ statistics above 200 were selected for testing. For genes that were positively correlated with capsule size, the gene deletion mutant was tested in inducing conditions (RPMI, 37°C, 5% CO_2_, no cAMP) with the prediction that deletion would reduce capsule size compared to WT grown in the same conditions. For negatively correlated genes, the gene deletion mutant was tested in almost-inducing conditions (RPMI, 37°, room air, no cAMP, no buffer) and non-inducing conditions (YPD, 37°, 5% CO_2_, with or without cAMP, no buffer).

### Forming metagenes as features for machine learning

From the original gene expression data matrix, we sought to decrease the number of features by filtering and combining genes. We first removed low variance genes by filtering out genes whose expression in >= 95% samples was within one log_2_ of their mean, indicating that less than 5% of samples showed substantial changes. We next combined highly correlated genes into metagenes (correlation threshold > 0.8). The “expression level” of a metagene was then set to the mean of the expression levels of its constituent genes. We repeated this process to combine genes into metagenes until we had combined all highly correlated genes into metagenes.

Strain verification by whole-genome sequencing

Mutant strains that showed an altered capsule phenotype were validated by whole-genome sequencing. We verified that these strains lacked the gene that was supposed to be deleted, had the expected selectable marker gene, and had no major structural variants. One strain from the Madhani collection which was expected to be isogenic to KN99 expect for deletion of CNAG_00328 turned out to also have a duplication of one half of chromosome 1. Because of this defect, we did not report its capsule phenotype in this paper.

1.	Longden, N.G. (Rhodes University, 1992).
